# Multi-run Concrete Autoencoder to Identify Prognostic lncRNAs for 12 Cancers

**DOI:** 10.1101/2021.08.01.454691

**Authors:** Abdullah Al Mamun, Raihanul Bari Tanvir, Masrur Sobhan, Kalai Mathee, Giri Narasimhan, Gregory E Holt, Ananda Mohan Mondal

**Author notes:** University of Miami, Coral Gables, FL 33146, U.S.A. Equal Contributors (Abdullah Al Mamun, Raihanul Bari Tanvir, Masrur Sobhan). Supplemental information (SI) is available at https://github.com/pwaabdullah/mrCAE.

## Abstract

Long non-coding RNA plays a vital role in changing the expression profiles of various target genes that leads to cancer development. Thus, identifying prognostic lncRNAs related to different cancers might help in developing cancer therapy. To discover the critical lncRNAs that can identify the origin of different cancers, we proposed to use the state-of-the-art deep learning algorithm Concrete Autoencoder (CAE) in an unsupervised setting, which efficiently identifies a subset of the most informative features. However, CAE does not identify reproducible features in different runs due to its stochastic nature. We proposed a multi-run CAE (mrCAE) to identify a stable set of features to address this issue. The assumption is that a feature appearing in multiple runs carries more meaningful information about the data under consideration. The genome-wide lncRNA expression profiles of 12 different types of cancers, a total of 4,768 samples available in The Cancer Genome Atlas (TCGA), were analyzed to discover the key lncRNAs. The lncRNAs identified by multiple runs of CAE were added to the final list of key lncRNAs, which are capable of identifying 12 different cancers. Our results showed that mrCAE performs better in feature selection than single-run CAE, standard autoencoder (AE), and other state-of-the-art feature selection techniques. This study discovered a set of top-ranking 128 lncRNAs that could identify the origin of 12 different cancers with an accuracy of 95%. Survival analysis showed that 76 of 128 lncRNAs have the prognostic capability in differentiating high- and low-risk groups of patients in different cancers. The proposed mrCAE outperformed AE, which select latent features and were thought to be the best tools for dimensionality reduction. By selecting actual features instead of pseudo-features, mrCAE can be valuable for precision medicine. The identified prognostic lncRNAs (this work), mRNAs, miRNAs, and DNA methylated genes (future work) can lead to biomarkers and therapies for different cancers.

## 1. Introduction

Recent studies showed that long non-coding RNAs (lncRNAs), which are longer than 200 nucleotides, play key roles in tumorigenesis (1–3). The lncRNAs also have key functions in transcriptional, post-transcriptional, and epigenetic gene regulation (4). Schmitt and Chang discussed the impact of lncRNA in cancer pathways (5). Hanahan and Weinberg described the involvement of lncRNAs in six hallmarks of cancer such as proliferation, growth suppression, motility, immortality, angiogenesis, and viability (6).

Hoadley *et al*. showed that cell of origin patterns dominate the molecular classification of tumors available in The Cancer Genome Atlas (TCGA) (7). Their analysis used copy number, mutation, DNA methylation, RPPA protein, mRNA, and miRNA expression. However, they did not consider another important molecular signature of cancer, which is lncRNA expression. This work motivated us to investigate the importance of lncRNAs in identifying different types of cancer. We hypothesize that there should be a shortlist of salient features or important lncRNAs with prognostic capability that could dictate the origin of multiple cancers.

In general, feature selection is worthwhile when the whole set of features is difficult to collect or expensive to generate (8). For example, in TCGA, the lncRNA expression profile dataset contains more than 12,000 features (lncRNAs) for each of 33 different cancers, and it is expensive to generate this data. Consequently, it is important to answer the question: *Is there a set of salient features (lncRNAs) capable of identifying the origin of 33 cancers or a subset of 33 cancers?*

Standard dimension reduction methods, such as principal component analysis (PCA) (9) and autoencoders (10) can generate a greatly reduced set of ***latent features***. However, these latent features are not ***the original features*** but are functional combinations of the original features. Identifying original features increases the “explainability” of the results and allows us to perform biological interpretation in diagnosing various deadly diseases, such as cancers. Recently, few deep learning-based feature selection methods showed improvement in selecting original features in both supervised and unsupervised settings (8,11–13).

In our previous study (14), we showed that a deep learning-based unsupervised feature selection algorithm CAE (8), performs better in feature selection, especially, in selecting a small number of features, compared to the state-of-the-art supervised feature selection methods such as LASSO, RF, and SVM-RFE. However, the study was based on the expression profiles of cancer patients only. The questions that remained unanswered are: (a) Were the identified lncRNAs cancer-specific or organ-specific? (b) CAE produces a different set of features in different runs, which raises questions about which set to use as the final feature set. (c) Do the identified lncRNAs have prognostic capability? (d) How to validate the identified lncRNAs?

In this paper, to address question (a), we analyzed expression data from 12 cancers identifying lncRNAs with fold change of at least 10. To address the question (b), we proposed to run CAE multiple times with a fixed number of features to be selected in each run and take the most frequently appearing features in multiple runs as the final set of features. To address question (c), survival analysis was performed to show that the identified features have prognostic capability. To address question (d), we checked the existence of identified lncRNAs in experimental works of literature, drug-lncRNA network, and cancer hallmarks.

The distribution of the number of samples for 12 cancers in TCGA is highly imbalanced, ranging from 36 for CHOL cancer to 1089 for BRCA cancer. Any supervised feature selection approach will be biased to heavy groups. So, the unsupervised nature of CAE can handle this issue in identifying appropriate features related to 12 different cancers. The proposed multi-run CAE approach filtered the key lncRNAs from 12,309 lncRNAs that are related to 12 different cancers with higher classification accuracy and better diagnosis of cancer origin compared to the state-of-the-art embedded feature selection approaches, including LASSO (15), RF (16), and SVM-RFE (17) and unsupervised feature selection approaches such as MCFS and UDFS.

Contributions of this study are as follows: 1) development of an optimal and stable feature selection framework, mrCAE. 2) Discovery of an optimal and stable set of 128 lncRNAs capable of identifying the origin of organs for 12 different cancers with an accuracy of 95%. 3) It has been shown that the lncRNAs identified using mrCAE from the expression profiles of cancer patients are truly cancer-specific, not organ-specific. 4) Survival or prognostic analysis of discovered lncRNAs. 5) Identified features, lncRNAs, are validated with existing literature, drug-lncRNA networks, and hallmark lncRNAs.

## 2. Materials and Methods

### 2.1. Data Preparation

To characterize the cancer-associated lncRNA, expression profiles and clinical data for 33 different cancers were downloaded from the UCSC Xena database (18). Each lncRNA expression was processed using a min-max normalization method to achieve good training performance. For this study, we considered the cancer types for which the number of normal samples is at least 10% of cancer samples, and 12 cancer types met this criterion. The distributions of cancer and normal samples for 12 cancers are shown in Table 1.

**Table 1.** Sample distributions of 12 cancers were considered in this experiment.

|  | BRCA | CHOL | COAD | KICH | KIRC | KIRP | LIHC | LUAD | LUSC | PRAD | READ | THCA |
| --- | --- | --- | --- | --- | --- | --- | --- | --- | --- | --- | --- | --- |
| Normal | 113 | 9 | 41 | 23 | 72 | 32 | 50 | 57 | 49 | 52 | 9 | 58 |
| Cancer | 1088 | 36 | 301 | 65 | 527 | 286 | 369 | 510 | 498 | 493 | 94 | 501 |

This dataset contains about 60 thousand RNAs expression profiles, including coding genes (mRNAs) and non-coding genes (lncRNAs and miRNAs). In this study, only the expression profiles of lncRNA (n = 12,309) were considered for analysis and model evaluation. The final dataset contains 4,768 cancer patients and 565 normal patients.

### 2.2. Feature Selection Using Multi-Run Concrete Autoencoder

For selecting important features (lncRNAs), a state-of-the-art deep learning-based unsupervised algorithm, Concrete Autoencoder (CAE) (8), was run multiple times iteratively. We name this approach multi-run CAE (mrCAE). The reason for using mrCAE is that CAE selects the most informative features in a *<u>stochastic manner</u>*, meaning different sets of informative features are selected in different runs. The **<u>assumption</u>** we made while running CAE multiple times is that if a feature appears in more than one run, that can be considered as a **<u>stable feature</u>**.

#### 2.2.1. Architecture and Working Principle of CAE

The architecture of a CAE (Figure 1) consists of a single encoding layer, also known as the “Feature selection layer” shown in yellow, and arbitrary decoding layers (e.g., a deep feedforward neural network), shown in the box on the right. The detailed algorithm is available in (8). The function of the encoder is to select a given number of *k* actual features (not latent features in the case of a traditional Autoencoder) in a *<u>stochastic manner</u>* from the original large input feature space, ***X*** of size *n*. The function of the decoder is to reconstruct the original features (***X***^′^ is the reconstructed feature vector) using the *k* features selected by the encoder.

**Figure 1:**
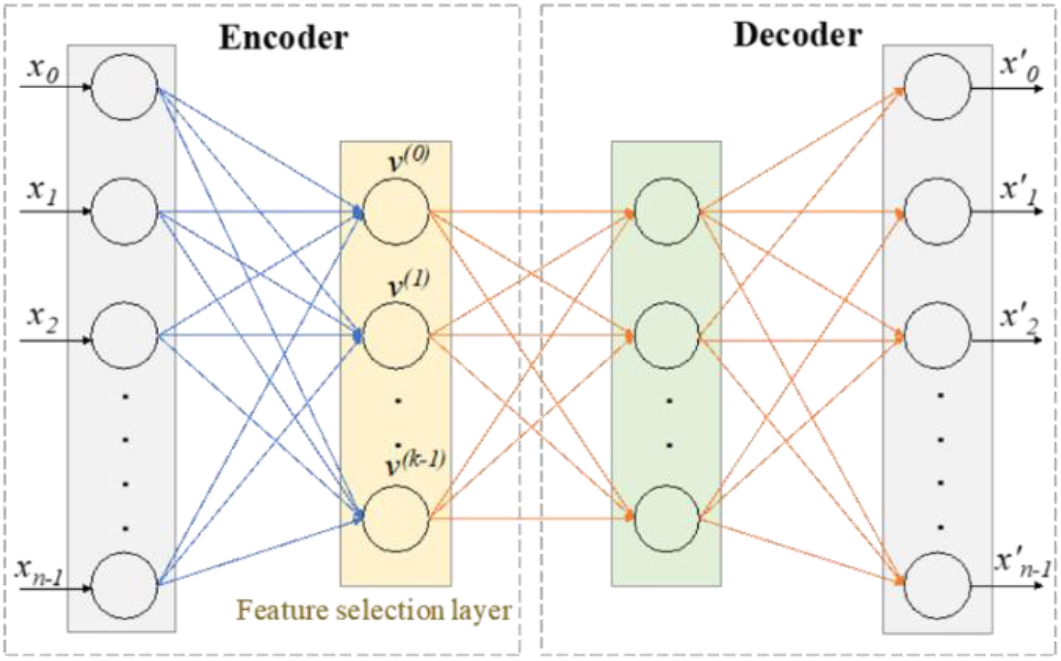
Architecture of Concrete Autoencoder. The architecture of CAE consists of an encoder and a decoder. The encoder is consist of an input layer and a concrete feature selection layer, shown in yellow. Feature selection layer has *k* number of nodes where each node is for a feature to be selected. Decoder is to check how well the input features can be reconstructed using the selected *k* features. Output layer has the same number of nodes as input layer. same as that of a standard autoencoder. *X* = [*x*_0_, *x*_1_, … *x*_*n*−1_] = Input features. 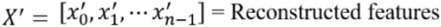.

How input features are selected depends on the *temperature* of the selection layer, which is modulated from a high value to a small value using a simple annealing schedule (8). As the temperature of the selection layer approaches zero, the layer selects *k* individual input features. The decoder of a concrete autoencoder serves as the reconstruction function. It is the same as that of a standard autoencoder. Thus, the concrete autoencoder is a method for selecting a discrete set of *k* features that are optimized for an arbitrarily complex reconstruction function.

### Training and Testing/Validation of CAE

The samples in a cohort are divided into 80/20 split for training and testing. In the training phase, 80% of samples are used to select the *k* informative features. In the testing/validation phase, 20% of samples are used to reconstruct their original features using the selected *k* features.

#### 2.2.2. Hyperparameter Tuning for CAE

The hyperparameters of CAE were tuned for lncRNA expression data of 12 TCGA cancer types. We kept two of the parameters the same as used in the original CAE, developed by Abid et al. (8). These two parameters are leaky ReLU with a threshold value of 0.1 and a 10% dropout rate. To tune the number of nodes in two hidden layers of the decoder, the model was tested by varying the number of nodes from 240 to 340, with a step size of 10. It was found that a decoder with 300 nodes in both layers yields the highest accuracy. So, the number of nodes in two hidden layers of the decoder was selected as 300.

To tune the number of epochs and learning rate, the random search (19) approach was used. For the number of epochs, the values used were 200, 300, 500, 1000, 1500, 2000, 2500, and 3000. Similarly, for learning rate, the values were 0.001, 0.002, 0.005, 0.0005, 0.01, and 0.05. In every run of CAE, the values of the two hyperparameters were randomly selected. With 300 epochs and 0.002 learning rate, the 100 features selected by the CAE produced the highest accuracy in classifying 12 cancer types using SVM. So, these parameter values were chosen for further analysis. Details of hyperparameter tuning are available in *Supplementary 1*.

For every iteration of a single run in the hyperparameter tuning phase, temperature, mean-max probability (mean of maximum probabilities of the selected features), training loss, and validation loss were observed and plotted. The plot paints a clear picture of the learning process in CAE at every epoch, thereby naming it as the characteristic plot of CAE, as shown in Figure 2.

**Figure 2.**
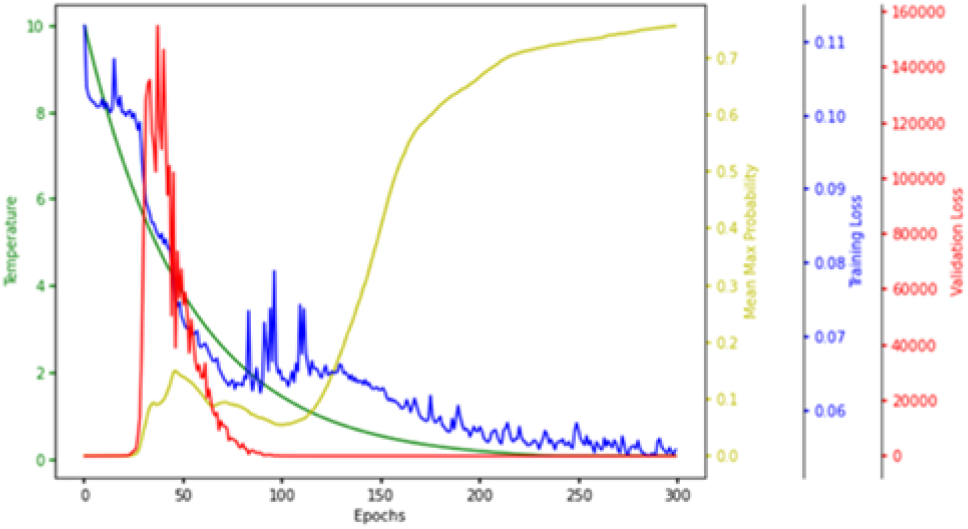
Characteristic Plot of Concrete Autoencoder. Temperature (green), mean-max probability (yellow), training loss (blue), and validation loss (red) are plotted at different scales.

One of the main objectives of this plot is to see if the model is converging in terms of loss, which is evident in Figure 2, as the training loss and validation loss converge to a lower value. Each node in the concrete selection layer learns a probability value for every feature, and the node selects the one with the highest probability. The higher the mean-max probability is, the more each node in the concrete selector is confident of one of the features. So, it is one of the goals to have the mean-max probability as high as possible.

### 2.3. Comparing mrCAE with Other Feature Selection Approaches

The feature selection capability of mrCAE was compared with the standard autoencoder (AE), three frequently used embedded feature selection models, including LASSO (15), Random Forest (RF) (16), and Support Vector Machine with Recursive Feature Elimination (SVM-RFE) (17), and two unsupervised feature selection models, MCFS (20) and UDFS (21). The exact number of features were selected, and those features were used to evaluate the classification performance in classifying 12 different cancer types. A stratified 5-fold cross-validation using SVM with linear kernel was conducted to evaluate the classification performance. Four different evaluation metrics - accuracy, precision, recall, f1 score - have been used to record the classification performance.

### 2.4. Implementation of Feature Selection Algorithms

All feature selection algorithm except mrCAE were implemented using the scikit-learn framework (https://scikit-learn.org/), whereas mrCAE was implemented using a deep learning framework named Keras (https://keras.io/). Experiments are parallelized on NVIDIA Quadro K620 GPU with 384 cores and 2GB memory devices. The dataset was split into the train and test set according to the 80/20 ratio to avoid overfitting. The training set was used to estimate the learning parameters, and the test set was used for performance evaluation.

## 3. Results and Discussion

The lncRNA expression profiles of 12 cancers were analyzed with the goal of identifying the key lncRNAs using mrCAE. ***First***, we showed that the features selected by CAE are truly cancer-specific, not organ-specific. ***Second***, we showed the stochastic nature of CAE in selecting equally significant different sets of features in different runs. ***Third***, we showed that mrCAE performed better than the single-run CAE and other state-of-the-art feature selection methods, including LASSO, RF, SVM-RFE, MCFS, and UDFS. ***Fourth***, we determined a stable set of lncRNAs that not only can stratify 12 different cancer types but also have the highest number of lncRNAs with prognostic behavior. ***Fifth***, optimal number of runs for mrCAE.

### 3.1. Features Selected from Tumor Tissues are Cancer-Specific, not Organ-Specific

To check that the features selected by CAE from the lncRNA expression profiles of cancer samples are truly cancer-specific, not organ-specific, we ran CAE separately on tumor and normal samples to identify two sets of 80 features (lncRNAs). Figure 3a shows only five commons between 80 tumor and 80 normal features, which evidenced that 75 out of 80 features are unique to both tumor and normal tissues. It is clear from the t-SNE plots of Figures 1b and 1c that tumor and normal features can distinctively cluster 12 tumor tissues and corresponding normal tissues, respectively. However, when we do the cross, meaning the t-SNE plot of tumor tissues using normal features (Figure 3d) and t-SNE plot of normal tissues using tumor features (Figure 3e), there are no distinct clusters for 12 tumor and corresponding normal tissues. *Supplementary 2* shows the similar results for 40-feature and 60-feature scenarios. These experiments proved that the features derived from tumor samples are truly cancer-specific, not organ-specific.

**Figure 3.**
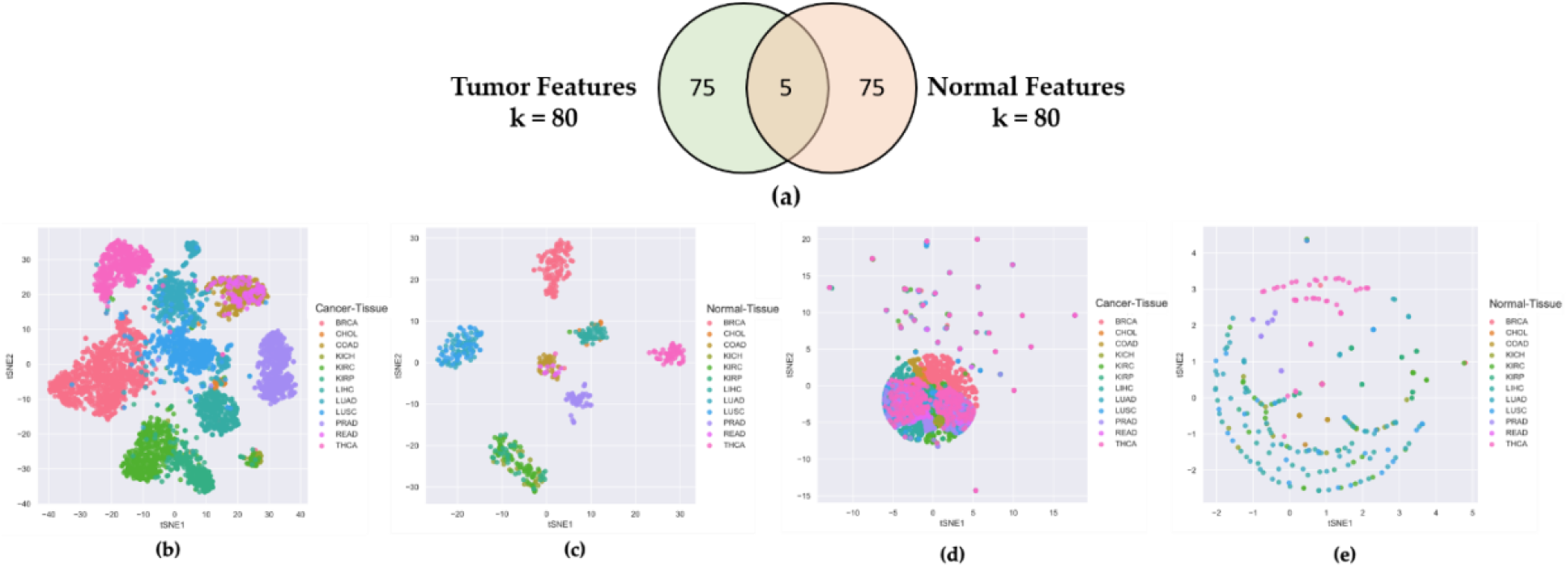
Comparing Tumor Features with Normal Features. a) Venn diagram of 80 tumor features and 80 normal features derived from CAE; b) t-SNE plot of tumor samples using tumor features; c) t-SNE plot of normal samples using normal features; d) t-SNE plot of tumor samples using normal features; e) t-SNE plot of normal samples using tumor features.

### 3.2. CAE Produces Different Sets of Significant Features in Different Runs

Though CAE selects a subset of the most significant features from a given dataset, it produces different sets of significant features in different runs due to its stochastic nature (12). To show the stochastic nature of CAE, three sets of 60 features were selected for the experiment. Figure 4 shows (a) the Venn diagram, (b) classification accuracy of 12 cancer types, (c) mean squared error (MSE) of reconstructing original feature space, and (d) t-SNE plots of clustering 12 different types of cancer samples, respectively, using three sets of features. It is clear from the Venn diagram that three runs selected three different sets of 60 features with slight inter-set overlaps of 10, 10, and 13 features. However, their performances in classifying 12 different cancer types are equally good (Figure 4b), as evidenced by the classification accuracies of ∼ 93%. Similarly, the three sets of features produced the same level of reconstruction errors of ∼ 0.06 (Figure 4c) and distinctive clusters of 12 different cancer types using t-SNE plots (Figure 4d).

**Figure 4.**
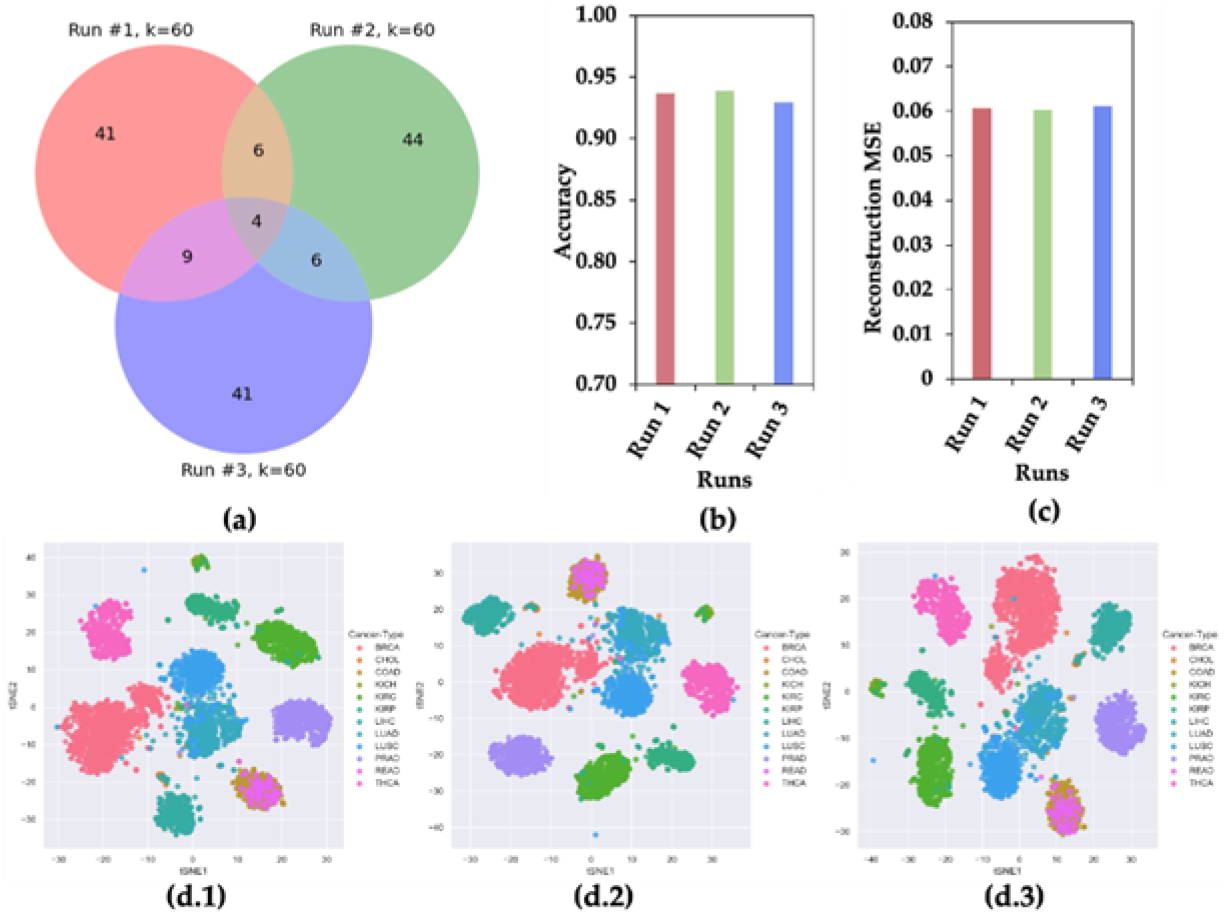
CAE Property of Selecting Different Sets of Features in Different Runs. a) Venn Diagram, b) accuracy of classifying 12 cancer types, (c) reconstruction mean squared error (MSE), and (d) t-SNE plots for 12 cancer samples using three sets of 60 features selected in three runs.

It can be concluded from Figure 4 that CAE selects different sets of most informative features in different runs. This observation motivated us to **hypothesize** that a feature appearing in multiple runs of CAE (mrCAE) carries the most meaningful information for a given dataset. The following section showed that mrCAE performed better than the single-run CAE and other state-of-the-art feature selection methods, including LASSO, RF, SVM-RFE, MCFS, and UDFS.

### 3.3. Comparison of mrCAE with Existing Feature Selection Approaches

Before comparing mrCAE with the existing feature selection approaches, we evaluated the performance of single-run CAE with a different number of selected features, which will guide us on how many features we should select for comparison. In Figure 5(a), it is noticeable that even with a smaller number of features, only with ten features, the average accuracy of CAE was close to 85%. There is a sharp increase in average accuracy (91%) with 20 features, followed by a slight increase (92% accuracy) up to 60 features. Then the curve reaches a plateau. From this figure, it seems like 40 features (before starting plateau) selected using different algorithms would be a good choice for comparison.

**Figure 5.**
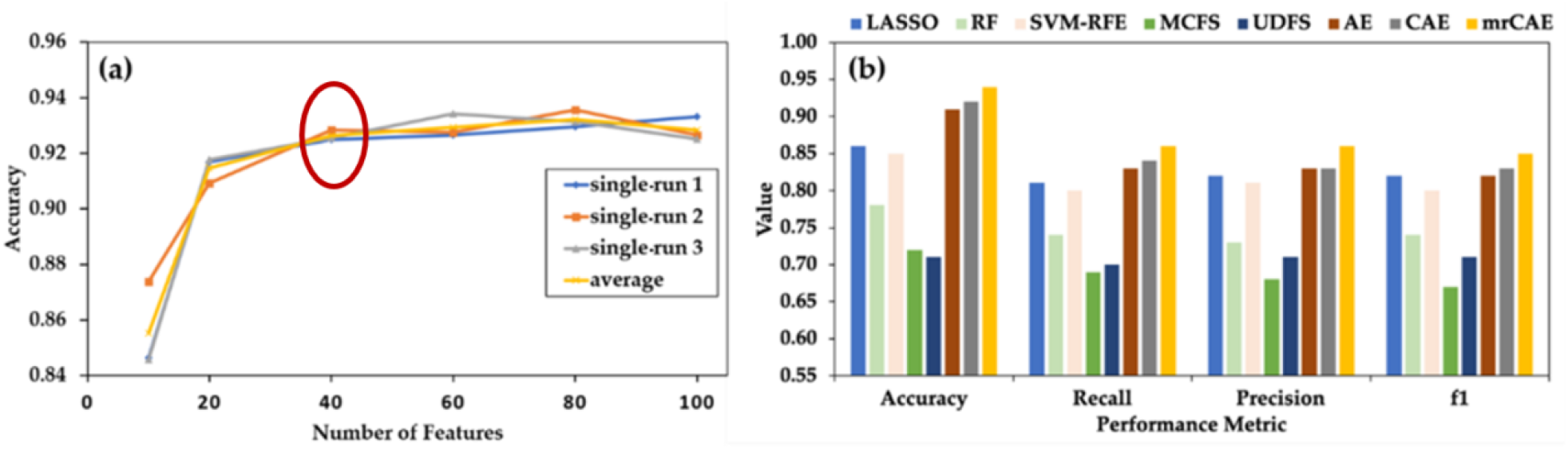
Comparing mrCAE with Ollier Feature Selection Approaches. **(a)** Behavior of single-run CAE to decide the number of features to be selected for comparison. CAE was run three times to select six sets of 10, 20, 40, 60, 80, and 100 features, “single avg” represents the average accuracy of three runs, **(b)** Classification Performance using forty features selected by LASSO, RF. SVM-RFE. MCFS, UDFS, AE, CAE, and mrCAE. Note that each approach selects 40 actual features except AE, which selects 40 latent features

#### Selection of 40 Features from mrCAE

CAE was run 100 times to select 100 features in each run. In 100 runs, it selected a total of 534 unique features. The frequency of appearance of these features in 100 runs ranged between 1 and 98. The top 40 features were chosen from the list sorted by decreasing frequency and were used to measure the performance of mrCAE.

Figure 5(b) shows the classification performance using the sets of 40 lncRNAs selected from different feature selection algorithms, including LASSO, RF, SVM-RFE, MCFS, UDFS, AE, CAE, and mrCAE. It is clear that mrCAE performed better than any other feature selection approaches regarding the accuracy, recall, precision, and F1 score.

### 3.4. mrCAE to Select a Stable Set of lncRNAs

#### mrCAE System

To identify a unique and stable set of lncRNAs that not only can distinguish between 12 different cancer types but also have the highest number of features with prognostic behavior, we designed mrCAE systems with 10, 20, 40, 60, 80, 100, 120 runs. 100 lncRNAs were selected in each of the single runs of an mrCAE system. Table 2 shows the summary statistics of mrCAE systems, including the total number of unique lncRNAs selected and the maximum frequency of a lncRNA appearing in each mrCAE system. The minimum frequency was 1 for all the different mrCAE systems. As shown in Table 2, a total of 223 unique lncRNAs (combined list of 10 sets of 100 lncRNAs) were selected by the 10-run mrCAE system, and the frequency of a lncRNA appearing in multiple runs ranges between 1 and 10. Similarly, a total of 575 unique lncRNAs were selected by the 120-run mrCAE system, and the frequency of a lncRNA appearing in multiple runs ranges between 1 and 117.

**Table 2.** Summary statistics of mrCAE systems in selecting lncRNAs.

| mrCAE | Total lncRNAs | Max Frequency |
| --- | --- | --- |
| 10-run mrCAE | 223 | 10 |
| 20-run mrCAE | 313 | 20 |
| 40-run mrCAE | 400 | 40 |
| 60-run mrCAE | 464 | 60 |
| 80-run mrCAE | 499 | 80 |
| 100-run mrCAE | 534 | 98 |
| 120-run mrCAE | 575 | 117 |

#### Frequent and Stable Features

Features appearing more than once in the mrCAE system are considered frequent features. Features with higher frequencies are considered stable features.

#### The Top Frequent Features

The top frequent features, for example, Top-10 features in any mrCAE system, are the first ten features from the combined list sorted in descending order based on frequency. To identify a stable set of lncRNAs, we selected the top features from each of the seven mrCAE systems in six different categories such as Top-10, Top-20, Top-40, Top-60, Top-80, and Top-100. Table 3 shows the ranges of frequency for the top features in six different categories. It is noticeable that the most frequent feature appeared in 10, 20, 40, 60, and 80 runs in the case of 10-, 20-, 40-, 60- and 80-run mrCAE systems, respectively, and the trend was not maintained for 100- and 120-run systems. In other words, the most frequent feature appeared in each run of each mrCAE system except the 100-run and 120-run systems, for which it (most frequent feature) appeared in 98 and 117 runs, respectively. It can be concluded that for the given lncRNA expression profile dataset of 12 cancers, an mrCAE system with 100 or more runs *<u>cannot produce the most frequent features in each run</u>*. Thus, 100-run mrCAE can be considered the optimal configuration for this dataset, and the results from 120-run mrCAE are not considered for subsequent analysis.

**Table 3.**
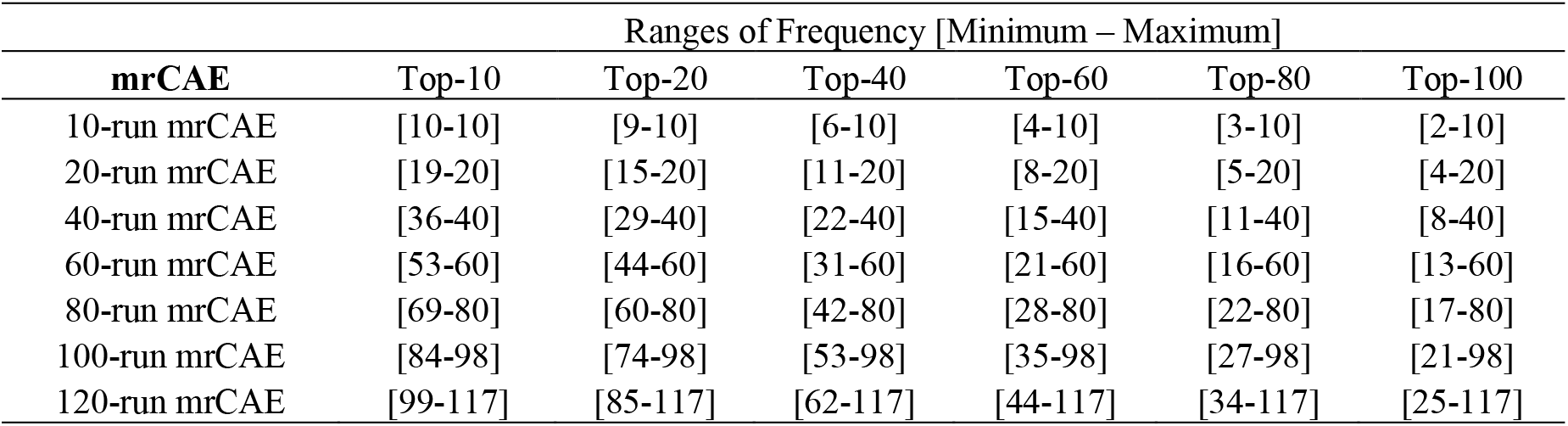
Ranges of frequency for the top features in six categories.

Finally, this experiment results in six unique sets of features corresponding to Top-10, Top-20, Top-40, Top-60, Top-80, and Top-100 features, as shown in the Venn diagram of Figure 6. For example, combining six sets of top-10 features from 10-, 20-, 40-, 60-, 80-, and 100-run mrCAE systems produces a unique list of 14 lncRNAs.

**Figure 6.**
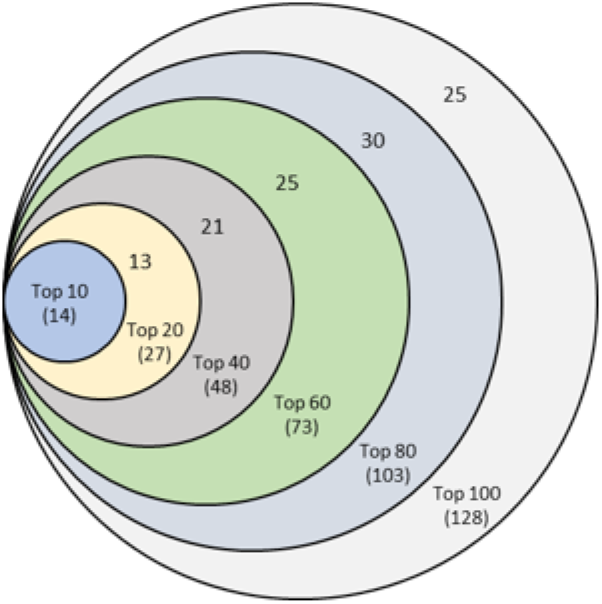
Venn diagram of six sets of unique features identified from six mrCAE systems. mrCAE consists of 10, 20, 40, 60, 80, and 100 runs. Each of these runs was conducted to select 100 features. The smallest set (light blue), containing 14 features, represents the unique feamres from six sets of Top-10 features from 10-, 20-, 40-, 60-, 80-, and 100-run mrCAE systems. Similarly, the 2nd smallest set contains 27 (14 + 13) unique features from six sets of Top-20 features and so on.

It is clear from the Venn diagram that each set of unique features is a subset of the following more extensive unique feature set. Finally, we can conclude that the 128 unique features (*Supplementary 3*), produced from the union of six sets of Top-100 features coming from 10-, 20-, 40-, 60-, 80-, and 100-run mrCAE systems, represent the stable and optimal feature set. We used this set of lncRNAs to do the downstream study, including survival and prognostic analyses and validation.

### 3.5. Prognostic Capability of Significant lncRNAs

To evaluate the prognostic capabilities of selected 128 stable lncRNAs, survival analyses on patients of different cancer types have been performed. Any lncRNA having zero expression values for most of the cancer samples was excluded from survival analysis of that cancer. The patients with values less than or equal to the median are labeled group A. Those with values greater than the median are labeled group B. After dividing into two groups, log-rank test was conducted, and the hazard ratio was calculated as the hazard rate of group A vs. hazard rate of group B to check the prognostic capability of a lncRNA. The criteria for a lncRNA to be prognostic are log-rank test P-value ≤ 0.05 and Hazard Ratio (HR) ≠ 1.0. Kaplan-Meier curve was plotted to show the prognostic behavior of a lncRNA.

Figure 7(a) shows the Kaplan-Meier plot for GATA3-AS1, one of the 11 prognostic lncRNAs for breast cancer, while Figure 7(b) shows the forest plot of survival analyses for 11 prognostic lncRNAs. It can be observed from Figure 7(a) that group B (red) has a higher rate of survival than group A(blue), meaning lncRNA GATA3-AS1 can successfully distinguish the high-risk group (Group A) of BRCA patients from the low-risk group (Group B). In other words, the cohort with low expression (blue) of GATA3-AS1 has 1.53 times higher rate of death compared to the high expression cohort (red). Thus, the cohorts with low expression values for seven lncRNAs (HR > 1.0) will have higher chances of death compare to high expression cohorts (Figure 7(b)). On the other hand, the cohorts with low expression values for four lncRNAs (HR < 1.0) will have lower chances of death compare to high expression cohorts (Figure 7(b)). *Supplementary 4* shows the forest plots for other cancer types.

**Figure 7:**
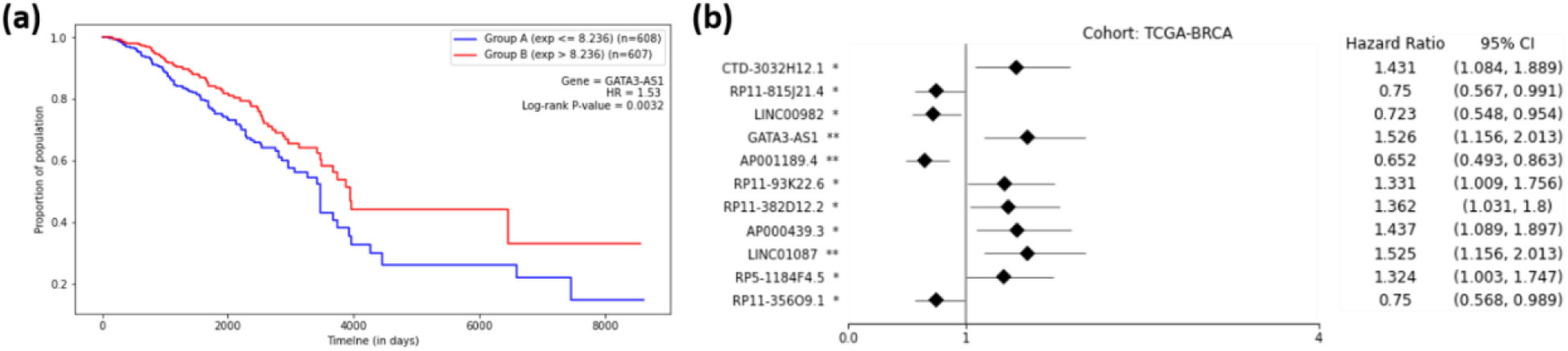
Survival Analysis of TCGA-BRCA. (a) Kaplan Meier Curve for lncRNA GATA3-AS1 on TCGA-BRCA cohort. Group A (blue) is the group with an expression less than or equal to the median, and Group B (red) is the group with an expression greater than the median. (b) Forest plot of survival analysis for 11 prognostic lncRNAs on BRCA cohort. The asterisks represent the Log-rank P-values (* - P ≤ 0.05, ** - P ≤ 0.01, *** - P ≤ 0.001, **** - P ≤ 0.0001).

The number of prognostically significant lncRNAs for each type of cancer is given in Table 4. The highest number of prognostic lncRNAs are discovered for KIRC (31 lncRNAs) followed by LUAD (22 lncRNAs), LUSC (18 lncRNAs). The proposed approach failed to discover any prognostic lncRNA for CHOL. The reason could be that the cohort consists of only 36 patients (Table 1). Some of the lncRNAs are prognostic for more than one cancer. Of 128 stable set of lncRNAs, 76 are prognostic.

**Table 4.** Summary of survival analysis regarding the number of prognostic lncRNAs for each of the 12 TCGA cancer types.

| BRCA | CHOL | COAD | KICH | KIRC | KIRP | LIHC | LUAD | LUSC | PRAD | READ | THCA | Total |
| --- | --- | --- | --- | --- | --- | --- | --- | --- | --- | --- | --- | --- |
| 11 | 0 | 3 | 3 | 31 | 15 | 1 | 22 | 18 | 4 | 4 | 10 | 76 |

### 3.6. Validations

The stable set of 128 lncRNAs derived from mrCAE were validated using existing literature (22). Of 128 lncRNAs, 103 are known lncRNAs (*Supplementary 5*) associated with different cancer types, Figure 8(a). For example, 98 lncRNAs are associated with BRCA, 52 lncRNAs are related to LUAD, and 37 lncRNAs are related to KIRP. Some lncRNAs are also found in four different cancer hallmarks, Figure 8(b) (*Supplementary 6*); for example, six lncRNAs are related to cancer “Prognosis.” We also validated the top 128 lncRNAs with existing drug-lncRNA networks (*Supplementary 7*). We found that 113 out of 128 lncRNAs are associated with 24 different drugs primarily used in cancer-related treatments, as shown in Figure 8(c &d). For example, the drug “Nilotinib” is mainly used to treat a specific type of blood cancer, associated with 18 different lncRNAs (Figure 8(e). Drug-lncRNA network formed based on the Spearman correlation coefficient between lncRNA expression levels and the IC50 values of the drug (23).

**Figure 8.**
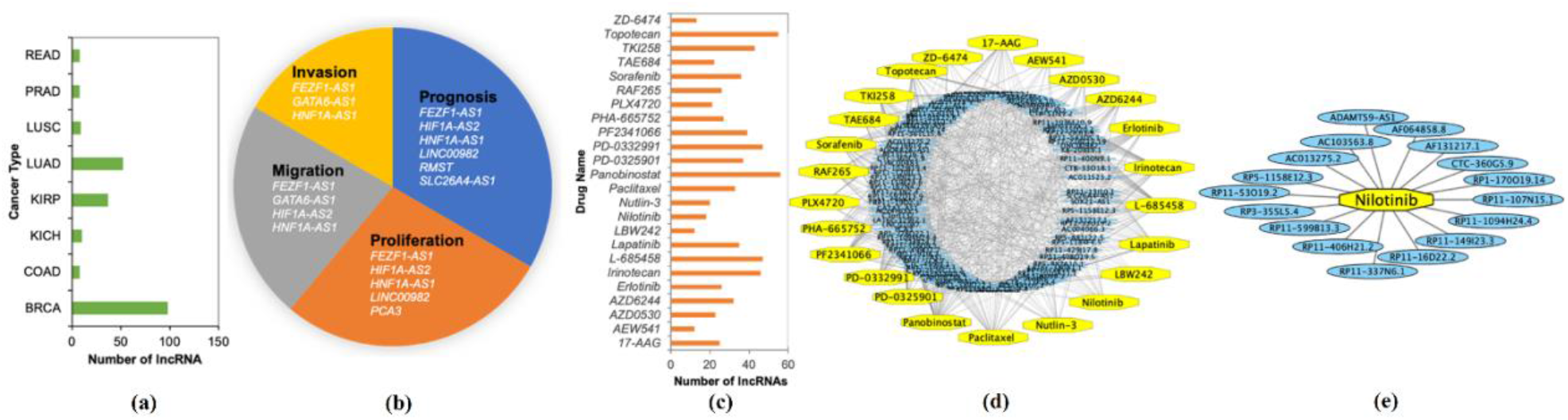
Validation of Identified lncRNAs. **a)** Number of known lncRNAs derived by mrCAE related to different cancer types found in (24–27) **b)** mrCAE derived lncRNAs related to different cancer Hallmarks (28). c) Number of lncRNAs related to different cancer drugs (23). d) Drug-lncRNA networks for all 24 drugs, e) An example lncRNAs-drug network for “Nilotinib”, which is used to treat certain blood cancers associated with 18 different lncRNAs.

## 4. Conclusion

In this study, we proposed a multi-run concrete autoencoder (mrCAE) to identify prognostic lncRNAs for multiple cancers. We tested the proposed model in analyzing lncRNA expression profiles of 12 cancers. The model selected a stable set of lncRNAs that can differentiate 12 cancers with high accuracy and provide subsets of prognostic lncRNAs for 12 cancers. Though the proposed mrCAE model is applied to multiple cancers, it can be used for single cancer, as used in identifying informative features for single-digit MNIST data by the developer of CAE, which will be our future work.

The lncRNAs selected by the proposed mrCAE outperformed the lncRNAs selected by the single-run CAE and other feature selection approaches. Also, the proposed mrCAE outperformed the standard autoencoder, which selects the latent features and is thought to be the upper limit in dimension reduction. Since the proposed mrCAE selects actual features and outperformed AE, it can provide information that can be used for precision medicine, such as identifying prognostic lncRNAs for different cancers.

## 5. Acknowledgments

This work has been partially supported by the NSF CAREER grant: #1651917 (transferred to #1901628) to AMM.

